# Alpha and beta desynchronization during consolidation of newly learned words

**DOI:** 10.1101/2025.01.09.632010

**Authors:** A. Zappa, P. León-Cabrera, N. Ramos-Escobar, M. Laine, A. Rodriguez-Fornells, C. François

## Abstract

While a growing body of literature exists on initial word-to-meaning mapping and retrieval of fully lexicalized words, our understanding on the consolidation that occurs between these two stages remains limited. The current study investigated the neural correlates of retrieving newly learned word meanings using oscillatory brain dynamics. Participants learned to associate new words with unknown objects and performed overt and covert naming tasks during the first and last days of a five-day training period. Behavioral results showed improved overt naming on Day 5 compared to Day 1. Selecting only words that were successfully produced in the overt naming task, we examined oscillatory activity associated with word retrieval while participants produced new words covertly, both pre- (Day 1) and post (Day 5) learning. The results showed a robust alpha (8-12 Hz) and lower beta (13-25 Hz) power decrease during covert naming after learning. We hypothesize that this alpha-beta power decrease indexes successful word retrieval following consolidation.

## 1. Introduction

Associating novel verbal labels to referents is a core learning mechanism essential for language acquisition. Following initial word-referent mapping, new lexical candidates eventually become integrated into existing semantic networks, allowing us to retrieve words from memory during language comprehension and production. Several studies have shown rapid cortical changes after very little exposure to lexical items in sentential contexts (Batterink & Neville, 2011; Borovsky et al., 2010; Lemhófer et al., 2025; Mestres-Missé et al., 2007; Perfetti et al., 2005; Shtyrov et al., 2010), which is sometimes interpreted as indexing new lexical entries (Gaskell & Dumay, 2003). However, it is a matter of debate whether these words have entered learners’ lexical-semantic network or whether they are temporarily associated with known concepts (for different perspectives, see Tamminen & Gaskell, 2013 or Rodriguez-Fornells et al., 2009). Indeed, it has been argued that lexicalization, or the integration of new items into a speaker’s existing lexicon, only occurs after a period of consolidation (Bakker et al., 2015a; Davis & Gaskell, 2009; Kaczer et al., 2018; Korochkina et al., 2024; Hulme & Rodd, 2023; Liu & van Hell, 2020; Schimke et al., 2022; Tamminen & Gaskell, 2013 but see Mestres-Misse et al., 2007 for a different perspective). Although a growing number of studies have investigated initial word encoding and the semantic processing of fully integrated words, the semantic processes unfolding between early word-to-meaning mapping and full lexicalization are still not fully understood.

The current study targets this intermediate phase, during which lexical candidates are progressively stabilized and integrated into long-term memory via consolidation mechanisms. According to the Complementary Learning Systems (CLS) framework (Davis & Gaskell, 2009; Gore et al., 20221; Singh et al., 2022), new information is initially encoded rapidly via the hippocampal system but only becomes fully integrated into neocortical memory networks over time, typically through offline consolidation during sleep. This gradual shift allows newly learned lexical items to be accessed more efficiently during comprehension and production. By investigating the retrieval of newly learned words after a multi-day learning period, the present study seeks to illuminate the neurophysiological mechanisms that support this transition toward stable lexical-semantic integration.

A number of previous studies investigating word learning have employed electroencephalography (EEG), a method that records electrical activity from the scalp and captures both time-locked responses to stimuli, known as event-related potentials (ERPs), and ongoing oscillatory activity across different frequency bands. Much of this work has focused the N400 ERP component as a neural marker of novel word learning (Bakker et al., 2015a; Batterink & Neville, 2011; Borovsky et al., 2010; Davis & Gaskell, 2009; Dittinger et al., 2016; Elmer et al., 2022; McLaughlin et al., 2017; Mestres-Missé et al., 2007). For instance, McLaughlin et al. (2004) observed a progressive emergence of the N400 to semantically related L2 word pairs over the course of a year-long French course, reflecting growing semantic integration. Other studies have demonstrated that N400 modulations can emerge after even brief exposures, with novel word forms acquiring semantic associations through contextual learning (Borovsky et al., 2010; Mestres-Missé et al., 2007; Batterink & Neville, 2011). Word learning has also been shown to engage predictive mechanisms, whereby pictures can pre-activate associated word forms. In a recent ERP study, N400 amplitudes to predictive pictures decreased as learning progressed, highlighting the N400’s sensitivity to emerging form-based associations (Elmer et al., 2022). Finally, a few studies have examined the emergence of semantic ERP effects following a period of consolidation after associative word learning (Davis & Gaskell, 2009), with mixed findings: some reported robust post-sleep semantic integration (Bakker et al., 2015a), while others failed to observe consistent effects (Takashima et al., 2019).

Importantly, the topography of the learning-related N400 varies across studies. For instance, differences in N400 spatial distribution have been associated with learning performance (Elmer et al., 2021) and cognitive control vs more automatic processing (Rodriguez-Fornells et al., 2009). The original study for which the current dataset was collected tracked changes in lexical-semantic processing across five training sessions via N400 modulations (Ramos-Escobar et al., 2021). In the first experiment, learners exhibited a frontal N400 during the initial stages of learning, which gradually decreased in amplitude and shifted to parietal sites during later stages. A second experiment teased apart learning and repetition effects by separating participants into a learning group and a non-learning control group, and replicated the initial findings in the learning group only. The change in N400 topography was interpreted as indicating that word learning requires both domain-general and language-specific mechanisms and involves distinguishable neural generators, depending on the learning stage.

The current study delved deeper into the neural correlates of consolidating new lexical items into memory. We used oscillatory brain dynamics, a decrease or increase in the power of neural activity in different frequency bands, thought to subserve cognitive processes as measured by EEG and magnetoencephalography (MEG), and more specifically event-related spectral perturbation (ERSP). Importantly, whereas ERPs only show phase-locked activity, ERSP shows both phase-locked and non-phase locked activity (Pfurtscheller & Silva, 1999, for a review see Hobson & Bishop, 2016) and could shed light on how lexical items are retrieved from memory during the early phases of word learning. A number of studies have examined word learning using brain oscillations and resting-state measures. Both Prat and colleagues (2016, 2019) and Huang and colleagues (2022, 2023) reported that individual differences in resting-state alpha and beta oscillations predicted success in artificial language learning. Küssner and colleagues (2016) also found that resting-state beta power predicted performance in a foreign vocabulary learning task. Kliesch et al. (2021, 2022) and Elmer et al. (2023) further explored the role of frequency-specific oscillatory dynamics and connectivity patterns in supporting word learning and memory consolidation. As a whole, these studies suggest that oscillatory activity during both resting state and learning tasks can provide valuable insights into neurocognitive readiness and plasticity during language acquisition. While the ERP and oscillation-based studies described above have been instrumental in identifying early neural markers of learning, our study shifts the focus to the outcomes of learning by investigating how newly formed word–referent associations are accessed during retrieval as understood in the prediction coding framework.

A growing body of literature relates alpha (8-12 Hz) and beta (13-25 Hz) power decreases to the encoding and retrieval of semantic representations (Branzi et al., 2023; Gastaldon et al., 2020; Griffiths et al., 2019; Hanslmayr et al., 2009, 2012; Khader & Rösler, 2011; Klimesch et al., 1999; León-Cabrera et al., 2022; Piai et al., 2014, 2015; Piai & Zheng, 2019). The magnitude of alpha and beta desynchronization has been shown to correlate with the amount of information retrieved from memory (Khader & Rösler, 2011) and is associated with greater semantic encoding and elaboration (Hanslmayr et al., 2009; Klimesch et al., 1999). This has brought about a comprehensive interpretation of alpha-beta desynchronization as reflecting the richness of the information represented in the system during semantic encoding and retrieval (Hanslmayr, Staudilg & Fellner, 2012; Griffiths et al., 2019). Within the framework of predictive language processing, desynchronization in these bands precedes contextually predictable words (Gastaldon et al., 2020; Momsen & Abel, 2022; Piai et al., 2014, 2015; Piai & Zheng; Wang, Hagoort, & Jensen, 2018). Indeed, recent studies in language comprehension have linked alpha and beta power decreases to the pre-activation (or, put differently, the anticipated retrieval) of the lexical-semantic features of final words in strongly constraining sentences, compared to weakly constraining ones (León-Cabrera et al., 2022; Li et al., 2017; Rommers et al., 2017; Terporten et al., 2019; Wang, Hagoort & Jensen, 2018). This pattern of oscillatory activity has been noted in both written and spoken language comprehension studies (León-Cabrera et al., 2022) and both for comprehension and production (Gastaldon et al., 2020). In short, alpha and beta decreases are widely interpreted as markers of semantic memory retrieval (Hanslmayr et al., 2012; Klimesch, 1999), including lexical-semantic retrieval (Branzi et al., 2023; Piai et al., 2015; Piai & Zheng, 2019; Hubbard & Federmeier, 2004). However, inconsistencies remain as to whether beta (Bakker et al., 2015b; Bastiaansen et al., 2005; Momsen et al., 2002), alpha (León-Cabrera et al., 2022; Strauß et al., 2014; Lago et al., 2023) or both frequency bands (Gastaldon et al., 2020; Hustá et al., 2021; Klimesch et al., 2001; Momsen & Abel, 2022; Piai et al., 2015) underlie these semantic-related processes. During the task used in this study, participants viewed pictures and were asked to name them covertly. We were particularly interested in the neural signature of the word retrieval process elicited by picture viewing.

Although the present study centers on alpha and beta band activity, previous studies have also implicated theta oscillations (4 -7 Hz) in the encoding and retrieval of verbal information. Increases in theta power have been linked to successful memory formation (Klimesch et al., 1996; Caplan & Glaholt, 2007; Hanslmayr et al., 2009; Osipova et al., 2006) and interpreted as reflecting hippocampo-cortical interactions that support the integration of new lexical items (Klimesch, 1999; for a review see Nyhus & Curran, 2010). Theta synchronization is often associated with the encoding of novel information, particularly in tasks requiring active learning or contextual integration (Zion-Golumbic et al., 2010; Momsen & Abel, 2022), though its role during post-consolidation word retrieval remains less well understood.

Only one previous oscillatory study examined the lexicalization of newly learned words post-consolidation using a word-learning task. Bakker and colleagues (2015b) taught participants novel words based on their native language and tested them during three different stages of lexicalization: before (untrained new words), right after training (recently learned new words), and 24 hours after training (new words trained one day before). Interestingly, decreased lower beta band power (16-21 Hz) and increased theta power (4-8 Hz) was induced for new words trained one day before when compared to untrained or recently learned new words. However, this consolidation effect was less clear or non-existent for upper beta (21-28 Hz) and alpha oscillatory bands. Indeed, recently learned words showed a decrease in power in the upper beta band, similar to new words trained one day before, raising doubts about the exact role of beta oscillatory activity in the consolidation of newly learned words. Although they tried to avoid motor artifacts by making it unpredictable whether participants had to answer or not, their task required a motor response and may have elicited confounding preparatory beta motor effects (Alegre et al., 2004). Importantly for the present study, Bakker-Marshall et al. (2018) followed up on the previous EEG study using MEG recordings and failed to observe a decrease in power in beta associated with new word consolidation. The authors noted the need to further investigate the intriguing role of beta power as an index of new word consolidation. More recently, Momsen and Abel (2022) used a contextual learning paradigm that allowed to build up the meaning of new words from context progressively (based on Mestres-Missé et al., 2007). They observed a decrease in alpha (8-12 Hz) and beta power (14-20 Hz) preceding the last presentation of the new word, which they interpreted as reflecting successful identification of the meaning of this word. These findings align in a compelling way with the abovementioned studies linking alpha and beta desynchronization prior to the pre-activation of contextually predictable words (León-Cabrera et al., 2022; Rommers et al., 2017; Wang et al., 2017; Terporten et al., 2019; Li et al., 2017), most probably associated with retrieving lexical and/or semantic features in highly constrained contexts (León-Cabrera et al., 2022; Gastaldon et al., 2020; Hustá et al., 2021; Klimesch et al., 2001; Momsen & Abel, 2022; Piai et al., 2015).

To gain a further understanding of the neurophysiological mechanisms of new word retrieval after learning, we used a covert naming task that aimed to isolate language-related alpha-beta desynchronization from motor-related effects often associated with overt articulation (Pfurtscheller & Lopes da Silva, 1999; Alegre et al., 2004; Piai et al., 2020), allowing a clearer interpretation of oscillatory activity as reflecting word retrieval. We performed analyses on the word-learning dataset collected on the first and fifth day of the study described above (Ramos-Escobar et al., 2021; see Fig. 1 for the task design). Word learning was measured pre- and post-training by covert naming (Figure 1), a form of internal word production that requires word-form retrieval from corresponding semantic representations (Dell & O’Seaghdha, 1992; Schmitt et al., 2000; Levelt, 1989; Rodriguez-Fornells et al., 2002; Piai & Zheng, 2019; Schriefers et al., 1990). Based on the previous literature cited above, we hypothesized that if novel word forms had been integrated into learners’ lexicon on Day 5, this would be indexed by a decrease in alpha and beta power during covert naming as compared to Day 1 (pre-training) due to participants having fully or partially created associations between new object representations and potential new words.

**Figure 1.**
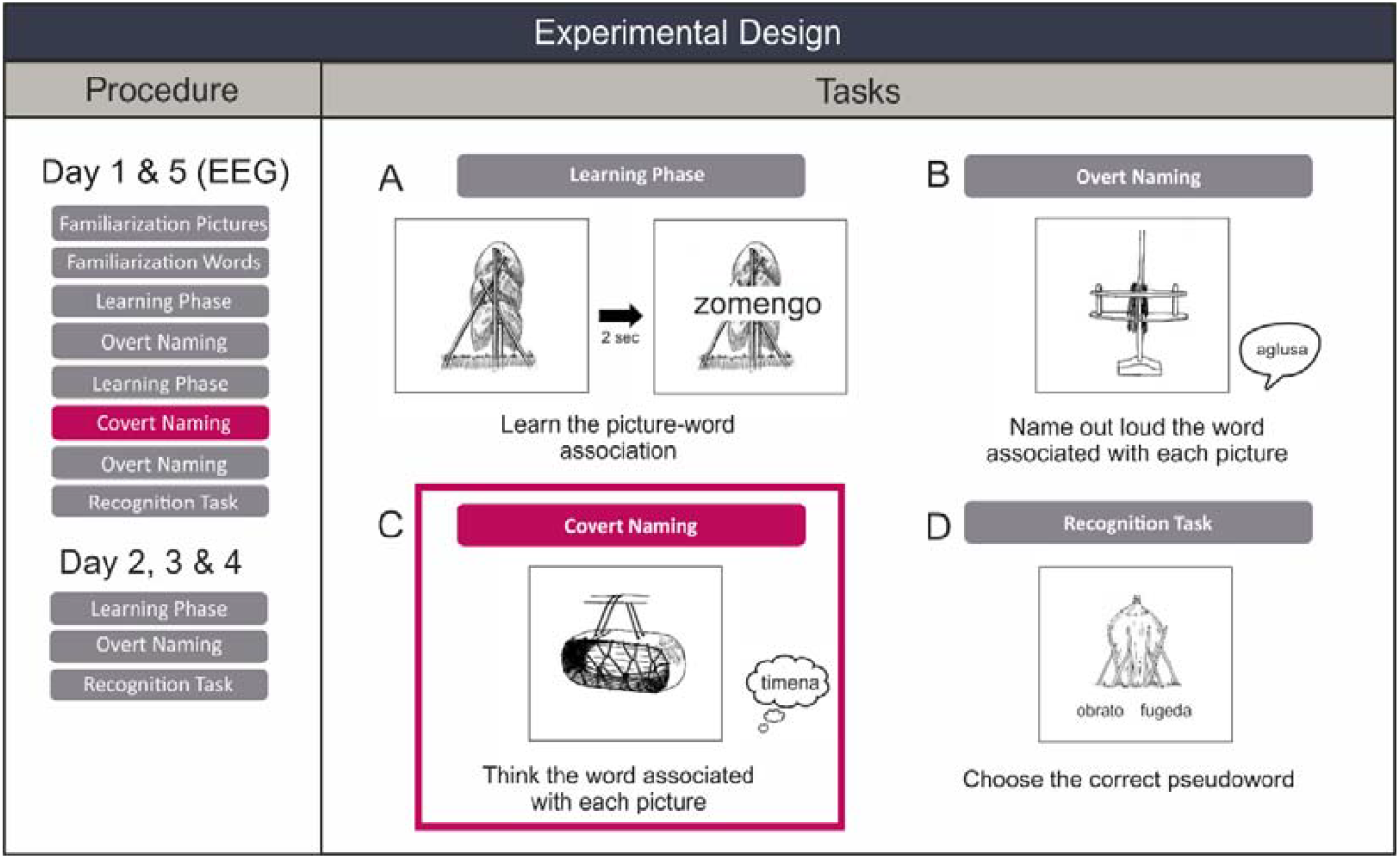
Adapted from the original study from Ramos-Escobar and colleagues (2021). Task design for the four different tasks used in the original study. B shows the the Overt naming task, from which we used the results to select the items that participants correctly named in our Covert naming analysis. Highlighted and framed in red is the Covert Naming task we focused on in the current study.

## 2. Methods

### 2.1. Participants

A total of 25 healthy volunteers (13 females, mean age: 22 ± 3.19 years) participated in the study. The data of three participants was lost due to technical problems during the covert naming task. In addition, since we were interested in tracking learning-related changes, we only kept the trials of items that were successfully learned by the participant (i.e., correctly named in the last overt naming task of Day 5), excluding those that were already learned on Day 1 (i.e., correctly named in the last overt naming task of Day 1) and thus not expected to undergo learning-related changes. Three participants were excluded because they had less than 35 correct trials available that fulfilled this criteria for analysis on Day 1 or Day 5. Therefore, 19 participants were included in the final data analyses (10 females, mean age = 21.8 ± 3.2 years).

Participants were all Spanish-Catalan bilinguals, except two Spanish monolinguals. They all had normal or corrected-to-normal vision, reported no history of neurological deficits, and were right-handed. Participants were briefed on the study procedure, provided written informed consent, and received a compensation of 60€ for their participation in the 5-day training sessions. The study received approval from the local ethics committee.

### 2.2. Procedure

The study consisted of five consecutive daily training sessions. On the first and last days, participants were first exposed to novel objects and new words during a pre-exposure task. Following this, they performed a learning task and an overt-naming task in the middle of learning. They then performed a covert-naming task, an overt-naming task, and a 2-alternative forced-choice (2-AFC) new word object matching task. On the second, third, and fourth days, participants engaged in the learning phase and the 2-AFC task. EEG data were recorded on Day 1 and Day 5 (**Figure 1**).

Before beginning the training sessions on the first day, participants completed a Language History Questionnaire (Bilingualism and Language Switching, Rodríguez-Fornells et al., 2012) and two tasks related to cognitive control and working memory: semantic and phonological fluency tasks (animal fluency and words starting with the letter ‘p’) and the WAIS-III Digit Span (mean score of forward and backward auditory span; see the original study for more details, Ramos-Escobar et al., 2021).

### 2.3. Stimuli and Task

A set of 139 unfamiliar black-and-white images depicting farming-related artifacts was selected from the Ancient Farming Equipment (AFE) word-learning paradigm (Laine & Salmelin, 2010). Twenty students from the University of Barcelona who did not participate in the present study rated these objects for familiarity (on a scale from 0, ‘totally unfamiliar,’ to 5, ‘very familiar’, mean rating: 2.58; SD: 0.61). Based on these ratings, the 120 least familiar objects were selected. A corresponding set of 120 tri-syllabic new words were created using the B-Pal software (Davis and Perea, 2005). These new words were phonotactically legal in Spanish and followed six different Consonant-Vowel (CV) structures (CVCVCV, VCVCVC, VCVCV, VCCVCV, CVCVCCV, and VCVCCV). They were presented visually.

#### Pre-exposure phase

Participants first completed a pre-exposure phase. In this stage, 120 novel objects were shown randomly for 2 seconds each. Additionally, five familiar objects from the AFE paradigm (those with the highest familiarity scores) were included. Participants were instructed to press a button when these target objects appeared to maintain a constant level of attention during this phase. This familiarization task (not reported here) was designed to examine brain responses to novel stimuli (both new words and images) when participants were instructed to observe rather than explicitly learn them. It was also intended to mitigate novelty-related ERP effects during the learning phase for the original study.

#### Learning phase

The 120 object-new word pairs were presented randomly in a blocked design. Participants were instructed to pay attention to object-new word associations and learn as many as possible. During learning trials, participants saw a novel object on the screen for 2 seconds. For the first second, the object appeared alone, and during the last second, the associated new word was displayed above it (**Figure 1A**). The initial part of the learning phase comprised 480 trials divided into four blocks, followed by a second part (after the overt naming task) of 240 trials organized into two blocks. To minimize fatigue, a short break was provided after every 30 trials.

#### Overt-naming task

This task was used to assess learning, testing learners’ ability to produce recently learned new words. Overt naming engages retrieval processes (Costa et al., 2009; Laine & Martin 2023), which facilitate learning when retrieval is suitably difficult (Agarwal et al., 2012; Pyc & Rawson, 2009; Roediger & Butler, 2011). 120 objects were displayed on the screen for 3 seconds, and participants were instructed to name the new word associated with the object overtly (**Figure 1B**). Feedback was not provided between trials. Eight randomized sequences of the images were created, with their presentation counterbalanced across sessions and participants to control for order effects. Behavioural responses were digitally recorded. Objects named without syllable or phoneme errors were considered correct. No mispronunciations were made as the new words respected the phonotactic rules of Spanish. However, when participants failed to pronounce a syllable or phoneme, the overt naming was considered incorrect. Participants completed two overt naming tasks on each day (Day 1 and Day 5). Only the behavioural (and not the EEG) results of the last overt naming task of each day are included in the current study.

#### Covert-naming task

The current study focused on this task, which was introduced before the last overt naming task. It was designed to induce active retrieval processes while recording EEG activity without the muscular artifacts that can occur during overt naming (Strijkers et al., 2011). As mentioned above, inducing retrieval under demanding conditions is also believed to facilitate learning, which was our second aim. Participants saw the 120 objects on the screen for 3 seconds and were instructed to think about the new word without overtly naming it (**Figure 1C**). A red square appeared around the object in 10% of the trials to maintain participants’ attention during the task. During these trials, participants were asked to name that object overtly. To avoid list effects, five randomized lists were created (one list for each time the task was administered) and counterbalanced across sessions and participants.

### 2.4. Behavioral data analysis

All responses in the overt naming tasks were transcribed and categorized as either correct or incorrect. Responses were only considered correct if the participant uttered exactly the same name. We computed the percentage of correct responses in the last overt naming task of Day 1 and Day 5 for every participant and subjected them to a dependent-samples t-test to assess differences in performance before and after the learning protocol.

### 2.5. EEG data acquisition and analyses

The EEG signal was recorded with tin electrodes mounted in an electrocap (Electro-Cap International) arranged in 29 standard positions (FP1/2, F3/4, C3/4, P3/4, F7/8, T3/4, T5/6, Fz, Cz, Pz, FC1/2, FC5/6, CP1/2, CP5/6, PO1/2, Oz) and using a BrainAmp amplifier (BrainVision acquisition software, Brain Products) with a sampling rate of 250 Hz. All electrode impedances were checked and kept below 5 kΩ during the recording session. The input signal was filtered with a high-pass filter at 0.01 Hz and a notch filter at 50 Hz to attenuate power line noise. The electrode in the FCz position served as the ground, an external electrode placed at the lateral outer canthus of the right eye was used as the online reference, and an electrode located in the infraorbital ridge of the right eye served to monitor vertical eye movements.

We analyzed the electrophysiological data recorded during the covert naming task on Day 1 and Day 5. The data were analyzed using the Fieldtrip toolbox version 20230108 (Oostenveld et al., 2011) running under MATLAB version 9.11 (R2021B). EEG data were re-referenced off-line to mean activity at the two mastoidal electrodes. For each participant and covert naming session (Day 1 and Day 5), the continuous EEG data were segmented into epochs of 3.1 s, encompassing 1.1 s before and 2 s after picture onset. The 10% of trials in the task where the participant had to name the object overtly were excluded from further analysis. Then, we performed artifact detection and rejection. First, we excluded epochs with voltage amplitudes ±100 microvolts in any electrode within the -0.1 to 1 s interval (time-locked picture onset). Additionally, any remaining epochs with artifacts (eye blinks, eye movements, electrode drifting, or muscle activity) were rejected through visual inspection. After artifact rejection, there were no statistically significant differences between the average number of trials on Day 1 (mean = 87.1 trials, SD = 10.4 trials) and Day 5 (mean = 87.8 trials, SD = 12.8 trials) [*t*(1,21) = –.263, *p* = .795].

As a result, and after applying the exclusion criteria (see the section on Participants above), the final sample (N = 19) had an average of 57 valid trials available per participant and session, i.e., on Day 1 or Day 5 (SD = 13.9 trials; min = 35 trials, max = 91 trials). There were no statistical differences in the number of available trials on Day 1 (mean = 56.7 trials; SD = 13.4 trials) and Day 5 (mean = 57.3 trials; SD = 14.7 trials) [*t*(1,18) = –.308, *p* = .761].

### 2.6. Time-frequency analyses

For every participant and session (Day 1 and Day 5), we computed the time-locked event-related potential and subtracted it from every trial-level time-locked data to keep only induced activity (Kalcher & Pfurtscheller, 1995; Rommers & Federmeier, 2017). Time-resolved power was computed by applying a Hanning taper with fixed length of 300 ms. This taper provided a spectral resolution of 3 Hz and temporal resolution of 300 ms, which was considered a good time-frequency trade-off to include a sufficient number of cycles considering the frequency bands of main interest (alpha/beta; alpha from 8-12 Hz and beta from 13-25 Hz approximately) and the probable onset and duration of the word retrieval processes (at or above 300 ms) that were at focus in the current study. Specifically, since we lacked an objective measure of naming onset, the expected time course of word retrieval was based on estimates suggesting that articulation for known objects can begin around 600 ms, and therefore word retrieval processes (i.e., conceptual preparation, lexical and phonological retrieval) are achieved within 200-600 ms post-picture onset (i.e., 400 ms average duration) (Indefrey, 2014). Accordingly, the taper was applied from 1 to 40 Hz in steps of 20 ms in the -1.1 to 2 s time-window, time-locked to the onset of the picture. The 1-second interval at the onset (−1 to -0.1 s) and offset (1 to 2 s) of the selected epoch served as a buffer to prevent edge artifacts. Frequencies below and above the alpha/beta band (from 1-7 Hz and above 25 Hz) were also decomposed to perform a data-driven, blind analysis, as described below.

For statistical analyses, the grand-averaged time-frequency data was baseline corrected to the 100 ms pre-stimulus period, and a cluster-based permutation analysis was applied on the -0.1 to 1-second interval time-locked to picture onset, including all electrodes and frequencies from 2 to 35 Hz. In brief, this statistical test identifies adjacent time points, electrodes and frequencies with similar differences between experimental conditions while successfully controlling for the family-wise error rate (FWER) (Maris & Oostenweld, 2007). Specifically, every sample (frequency x time x channel) was compared between the two sessions (Day 1 and Day 5) using a paired-sample-statistic. All samples that exceeded the t-value threshold of ± 2.10 (for an alpha level of 0.05 with 18 degrees of freedom) were selected and clustered on the basis of spectral, temporal and spatial adjacency. The t-values within every cluster were summed and the maximum cluster-level sum was used to compute the permutation p-value through the Monte Carlo method involving 5000 random permutations. Only effects with a permutation p-value below 5% (two-tailed testing) were considered significant.

## 3. Results

### 3.1. Behavioral results

For the overt naming task, we computed the percentage of correct responses for Day 1 and Day 5 and compared their means with a dependent samples t-test. There was a clear learning effect with participants increasing their number of correct responses between Day 1 (mean = 13.6%; SD = 11.3) and Day 5 (mean = 76.9%; SD = 18.9; t(1,18) = -18.6, *p* < .001, see **Figure 2**).

**Figure 2.**
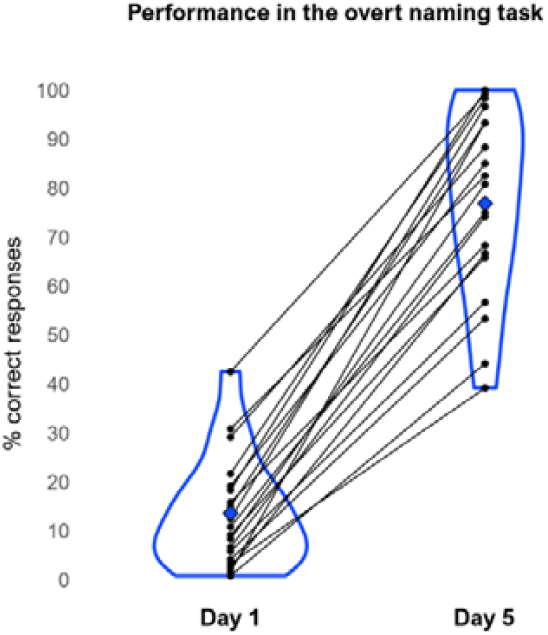
Violin plots of the performance (percentage of correct responses, y-axis) in the overt naming task on Day 1 and Day 5 (x-axis). The black dots represent individual participants and the lines connect data points from the same participant in each session. The blue diamond-shaped symbol indicates the group mean performance.

### 3.2. Time-frequency results

**Figure 3** shows the grand-averaged time-frequency maps of the covert naming task in each session (Day 1 and Day 5). As can be seen, picture presentation induced a long-lasting power decrease (strong desynchronization) of alpha (8-12 Hz) and lower beta (18-25 Hz) power after 150-200 ms on both days. This effect was predominant over posterior and occipital electrodes. In addition, there was a broadly distributed power increase in the theta band (4-8 Hz) approximately from 50-150 ms after picture presentation on both days.

**Figure 3.**
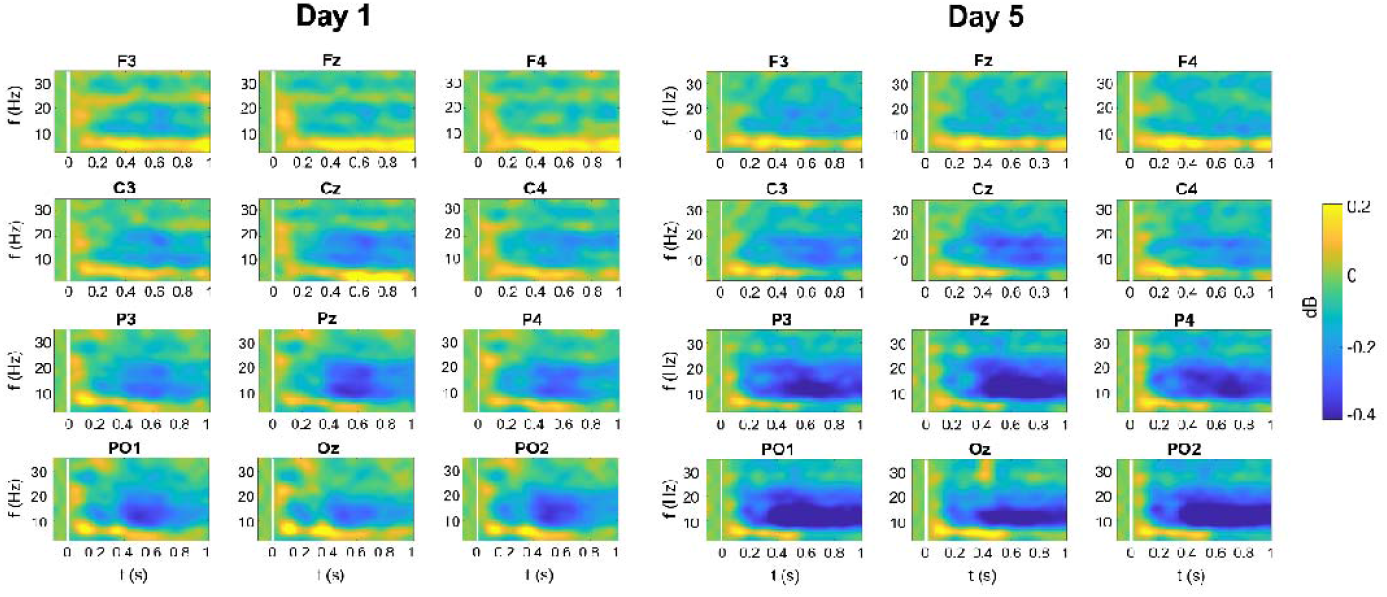
Grand-averaged time-frequency plots showing the oscillatory activity in the covert naming task of Day 1 and Day 5 (in dB) relative to baseline (−0.1 to 0 s time-locked to picture presentation) for frontal (F3, Fz, F4), central (C3, Cz, C4), posterior (P3, Pz, P4), and posterior-occipital (PO1, Oz, PO2) electrode positions. All plots encompass the frequency range from 0 to 35 Hz (y-axis) and the temporal interval from -0.1 to 1 second (x-axis) time-locked to picture onset. Picture onset is indicated by white vertical lines at 0 s.

Critically, the power decrease in the alpha and beta bands was more pronounced on Day 5 than on Day 1, suggesting that desynchronization in these frequency bands may reflect cognitive processes associated with the retrieval of the learnt new words. Accordingly, results of the cluster-based permutation test revealed one statistically significant negative cluster (*p* < .001), indicating a more pronounced power decrease on Day 5 compared to Day 1 (**Figure 4A**). The cluster mainly encompassed the alpha and lower beta bands and was most prominent at posterior and occipital sites within an interval ranging from about 150 ms to 1 second after picture presentation, although it also encompassed central and frontal electrodes (**Figure 4B**). In turn, the power changes in the theta band were not statistically significant between sessions.

**Figure 4.**
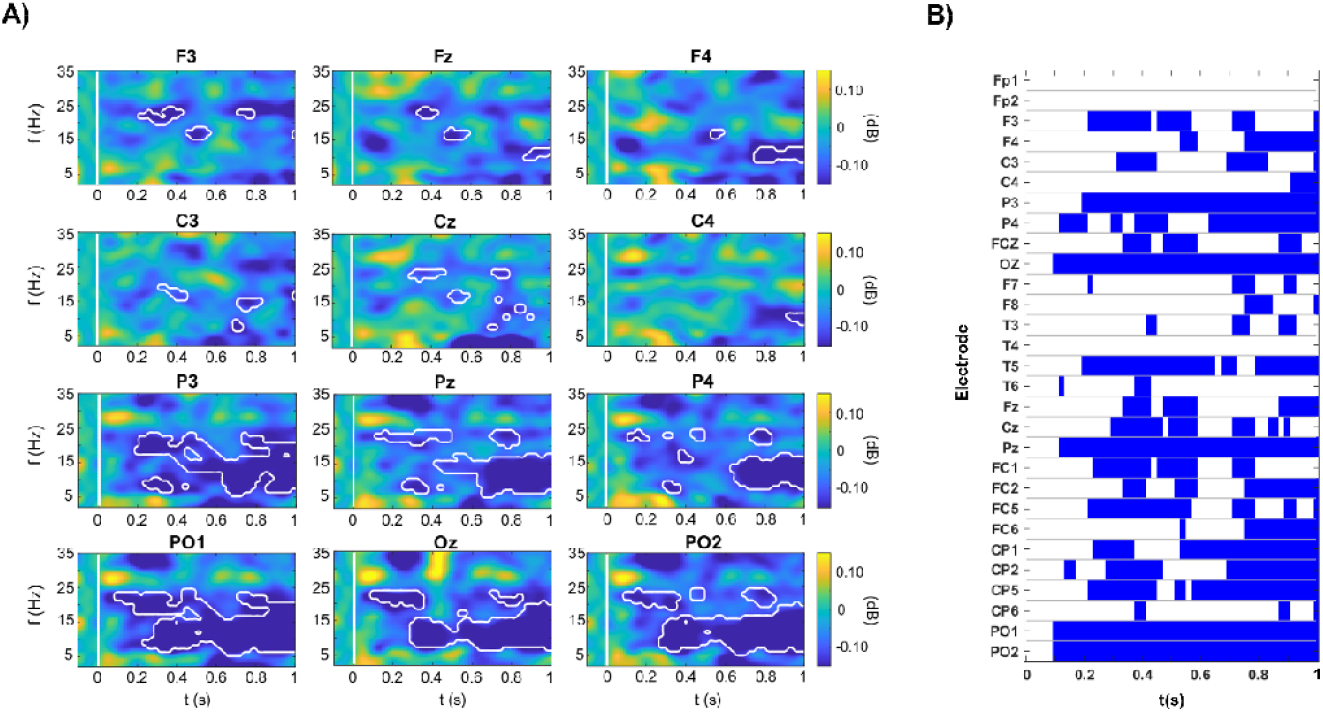
A) Grand-averaged time-frequency plots showing learning-related power changes (Day 5 minus Day 1) (in dB) relative to baseline (−0.1 to 0 ms time-locked to picture presentation). The statistically significant cluster is outlined in white. B) Raster plot of the time course of the significant cluster in the 8-25 Hz frequency range across all electrodes in the interval from 0 to 1 second after picture presentation.

Importantly, for each participant, we only kept the trials of words that were not learnt by Day 1 but that were eventually learnt on Day 5 (the learning status was evaluated through the performance in an overt naming task, see **Figure 2**). Therefore, it is likely that any difference between sessions reflects learning-related activity. Noteworthily, we performed an exploratory analysis on a smaller set of participants using incorrect trials. This additional analysis replicated the pattern of results and relevantly showed no evidence of reduced alpha-beta power on Day 5 on incorrect trials (i.e., items that were not produced or were incorrectly produced in the overt naming task; see **Supplementary Materials**). However, these results should be interpreted with caution given the small number of participants and trials involved in the analysis.

The finding of a single significant cluster supports an interpretation of concurrent alpha-beta desynchronization as a unitary phenomenon in this study. However, power decreases in these two frequency bands has also been associated with independent cognitive processes in the sensorimotor domain. For example, posterior-occipital power decreases in the alpha (but not in the beta) band have been associated with anticipatory attention (Foxe et al., 1998). Also, while both frequency bands support movement, they show spatial and functional dissociations (Brinkman et al., 2016; Stolk et al., 2019). Of special interest is the robust involvement of the beta band in motor-related processing (e.g., Salmelin & Sams, 2002), which is often a confound in production studies that seek to disassociate motor-related activity from semantic and lexical processing. Thus, to gain a deeper understanding of the cognitive processes involved in this task, we also inspected the spatial and temporal dynamics of these two frequency bands separately. **Figure 5** shows the evolution of the topographical distribution of mean power changes (Day 5 minus Day 1) at picture onset for the alpha and beta range individually. From 200 ms after picture onset, both frequency bands exhibited relatively enduring power decreases at parieto-occipital electrodes with a similar time-course (**Figure 6**). In addition, in the alpha band (8-12 Hz), right frontal electrodes exhibited a power decrease approximately from 400 to 1000 ms after picture onset. In turn, in the beta band (18-25 Hz), power desynchronized at left frontal sites from 200 ms to 1000 ms. Therefore, each frequency band might have contributed distinctly to the frontal effects captured by the cluster.

**Figure 5.**
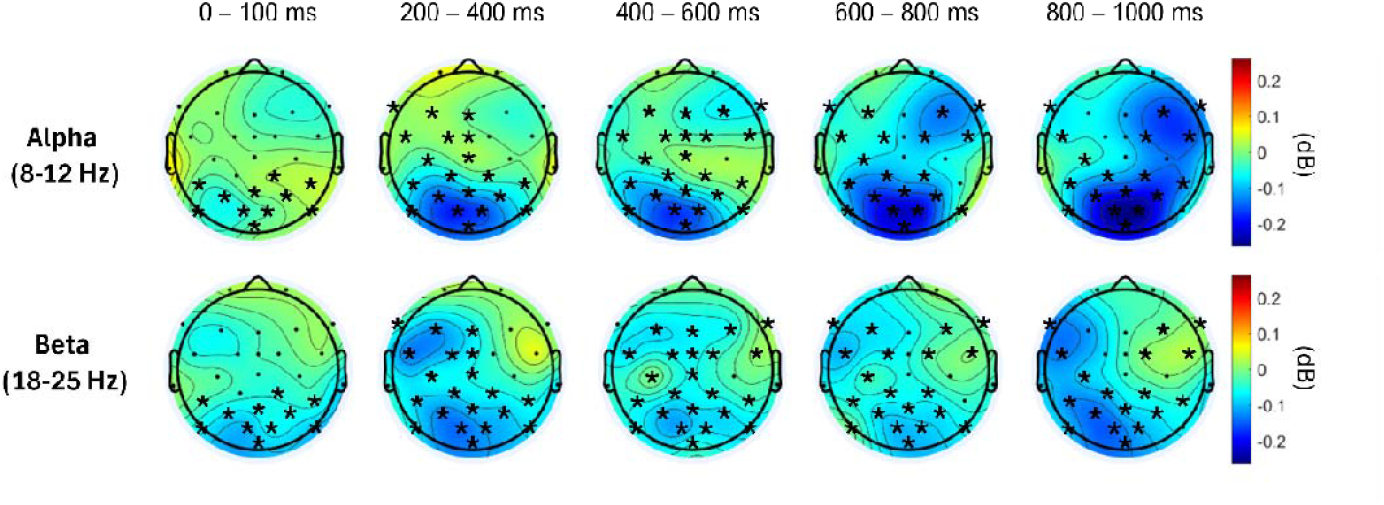
Topographical maps of the mean power changes (Day 5 minus Day 1; in dB) for the alpha (8-12 Hz) (top) and beta range (18-25 Hz) (bottom) in the 0 to 1 second interval time-locked to picture onset, in steps of 100 ms. In every time interval, asterisks mark the electrodes that were part of the s significant cluster which encompassed both frequency bands.

**Figure 6.**
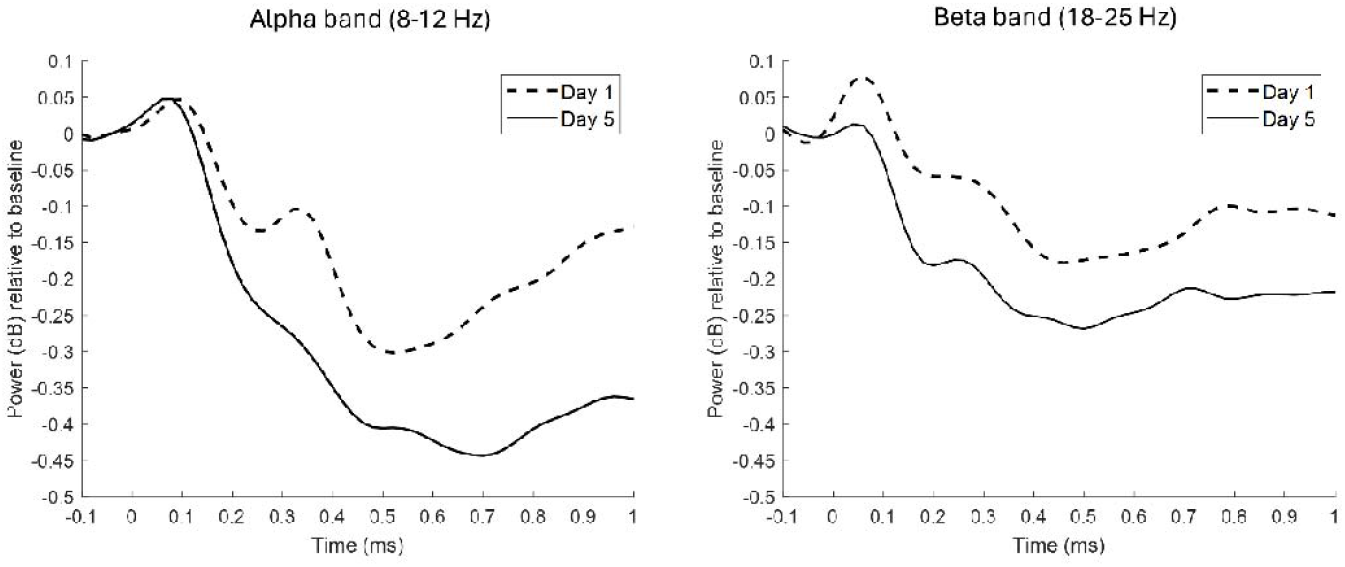
Time course of learning-related power changes in the alpha (8-12 Hz) (right) and lower beta (18-25 Hz) (left) bands for Day 1 (discontinuous lines) and Day 5 (continuous lines) at posterior-occipital electrodes (average of activity at Pz, P3, P4, PO1, PO2 and Oz electrodes).

## 4. Discussion

The current study explored the oscillatory activity in alpha and beta bands as participants retrieved newly learned words after consolidation. Previous studies provided conflicting results regarding the role of these frequency bands in new word learning (Bakker et al., 2015b; Bakker-Marshal et al., 2018; Momsen et al., 2022). Here, participants were taught to associate new words with unknown objects and asked to overtly and covertly name visually presented items during the first and last day of a five-day training experiment. We analyzed oscillatory brain activity related to retrieving new words focusing on alpha (8-12 Hz) and beta (13-25 Hz) band desynchronization while participants saw images and covertly retrieved new words, pre and post consolidation. We compared cortical oscillatory activity during covert naming at the first and fifth learning sessions, observing a robust concurrent alpha (8-12 Hz) and lower beta (18-25 Hz) power decrease 200-1000 ms post picture presentation. For both the alpha and beta frequency ranges, the effect was distributed over the parieto-occipital sites. When inspecting each frequency band separately, the power decrease in the alpha band encompassed left frontal electrodes from 400 to 1000 ms and the lower beta band (18-25 Hz) showed decreases in left frontal sites from 200 ms to 1000 ms.

One challenge in previous studies examining cortical oscillations during the retrieval of newly learned words is the potential confound introduced by motor responses (Bakker et al., 2015b). Alpha-beta desynchronization, while often linked to semantic processing, can also reflect visual processes (Hanslmayr et al., 2011) and cortical-subcortical motor activity (Neuper et al., 2006; Pfurtscheller & Lopes da Silva, 1999). Notably, in tasks requiring language articulation, alpha-beta desynchronization has been localized in the motor and premotor cortex and the left inferior frontal gyrus pre-speech, associated with verbal motor planning, even in the absence of semantic processing (Herman et al., 2013; Jenson et al., 2014). In the current study, we used a covert naming task that minimized overt motor activity, reducing the likelihood of motor-related confounds (Alegre et al., 2004; Pfurtscheller & Lopes da Silva, 1999). As such, we interpret the observed alpha and beta desynchronization as primarily reflecting word retrieval as opposed to motor activation or preparation (Hanslmayr et al., 2009). Nonetheless, we acknowledge that even covert naming may involve some degree of motor simulation or internal articulation.

To distinguish motor from language-related contributions of beta desynchronization, Scaltritti et al. (2020) compared silent reading and copy-typing tasks using emotionally valenced words. While motor-related beta desynchronization was specific to typing, both tasks elicited an earlier beta desynchronization linked to language processing. Similarly, Piai et al. (2020) used MEG to differentiate lexical-semantic retrieval from motor processes in tasks involving picture naming and conceptual judgment. For picture naming, beta desynchronization was localized to left temporal and ventral premotor areas, implicating lexical retrieval and verbal motor planning. By contrast, the judgment task elicited beta desynchronization in left posterior temporal and inferior parietal areas, alongside right motor cortex activation, associated with conceptual processing and manual response preparation. Another MEG study by Gastaldon et al. (2020) examined alpha-beta desynchronization during language comprehension and production in high- and low-constraint contexts. In comprehension tasks, highly constraining sentences elicited alpha-beta desynchronization in left lateralized language production areas, suggesting overlap between comprehension and production mechanisms. This activity also appeared over right posterior temporo-parietal regions, linked to internal modeling and contextual updating. Across these tasks, alpha-beta desynchronization indexed rich linguistic information encoding and retrieval, with its spatial distribution varying by task demands -whether lexical retrieval, language production, or motor preparation. Our findings show alpha and beta desynchronization in right frontal, left frontal and mainly parieto-occipital electrodes. Although EEG does not allow for specific localization claims, this does suggest that these effects did not occur in motor-related areas (i.e., ventral premotor areas as in Piai et al. 2020). This highlights the utility of alpha-beta desynchronization as a marker of language processing, distinct from motor contributions, and advances our understanding of the neurophysiological correlates of lexical-semantic retrieval.

As outlined in the introduction, alpha and beta desynchronization has been linked to the encoding and retrieval of semantic information, with greater power decreases reflecting richer or more elaborated lexical-semantic representations (Branzi et al., 2023; Gastaldon et al., 2020; Griffiths et al., 2019; Hanslmayr et al., 2009, 2012; Khader & Rösler, 2011; Klimesch et al., 1999; León-Cabrera et al., 2022; Piai et al., 2014, 2015; Piai & Zheng, 2019). Our results align particularly well with comprehension studies reporting alpha and beta power decreases over left frontal (Gastaldon et al., 2020; Wang et al., 2017; Rommers et al., 2017; León-Cabrera et al., 2022) and parieto-occipital electrodes (Rommers et al., 2017; León-Cabrera et al., 2022). Similarly, the alpha-beta desynchronization observed in this study might reflect lexical-semantic retrieval of newly learnt words that have been integrated into semantic memory after consolidation. This interpretation could also account for recent findings in other word learning studies linking alpha and/or beta to word learning (Bakker et al., 2015b; Momsel & Abel, 2022; but see Bakker-Marshal et al., 2018 for discrepant results). Although not directly related to word-learning, alpha power decreases have also been observed in a lexical decision task for real words versus ambiguous new words and for ambiguous new words versus full new words (Strauß et al., 2014).

Many previous studies investigating the neural correlates of retrieving newly learned novel words through oscillations have relied on lexical competition or semantic priming to measure word acquisition (Bakker et al., 2015b; Batterink & Neville, 2011; Havas et al., 2017; Hawkins & Rastle, 2016; Kaczer et al., 2018; Liu & van Hell, 2020; Tamminen & Gaskell, 2013). However, semantic priming does not necessarily indicate full lexicalization (see Korochkina et al. 2024 for a detailed discussion). In contrast, our study required participants to produce learned words overtly, and only those that could be explicitly recalled were included in the time-frequency analysis of covert naming. We believe this approach offers a more robust measure of novel word lexicalization, allowing for a clearer interpretation of the processes reflected in our time-frequency results. Similarly, in a recent study that used an incidental learning task, new words were learned through a dialogue-like situation and the final production of the new word was used as a potential measure of fast lexicalization (Lemhöfer et al., 2025).

Furthermore, a key distinction between our study and previous oscillatory ones (Bakker et al., 2015b; Momsen & Able, 2022) is our use of explicit word-to-referent associations to link new words to *unknown* objects – a task considered more demanding than associating novel words with familiar items or definitions (see Laine & Salmelin, 2010). In relation to this paradigm, it has recently been shown that learners tend to automatically assign meaning to unfamiliar referents quite rapidly after initial exposure (Laine et al., *in press*). Nonetheless, overnight consolidation likely plays an important role in processing information involving completely novel objects, as they cannot be readily associated with pre-existing familiar object representations (James et al., 2017; Schimke et al., 2021). Indeed, in a previous study, the pairing of new words to unfamiliar objects showed less beneficial sleep consolidation effects than when new words were associated with familiar objects (Havas et al., 2018). Thus, the capacity to use existing knowledge schemas to encode new representations might enhance the potential consolidation of the new traces. In the present study, new associative links are needed to be established between the new lexical candidate and the new unfamiliar object, as well as potential associations between this new object with already existing semantic information and other words in lexical-semantical networks (e.g., “*this new object resembles a tool I know*”) (Laine et al., *in press*). It is reasonable to conclude that during word learning, learners access and retrieve newly formed and progressively consolidated lexical-semantic representations that facilitate optimal information processing. This is in line with the CLS model (Davis & Gaskell, 2009), whereby new word representations become stabilized and integrated into existing networks during sleep (Gilboa & Moscovitch, 2021). Therefore, in our study, the observed alpha-beta desynchronization during the retrieval of newly learned words likely reflects the successful consolidation of these words.

Finally, several limitations must be considered. Firstly, the experimental design did not enable a comparison between learned and unlearnt items, which would have served as an optimal control for the contribution of stimulus repetition in the observed effects. Of note, we replicated the same pattern of alpha-beta power desynchronization while at least partially controlling for stimulus repetition effects (see **Supplementary Materials**). However, to appropriately control for this confound, future studies on covert naming could incorporate a within-subject manipulation that includes items that cannot be learned (i.e., random associations) (Elmer et al., 2021, 2022). Secondly, we lacked an objective estimate of the onset of word retrieval processes in the covert naming task. Instead, we based the decision to analyze the 1-second interval post-picture onset on the time course of word production stages, whereby conceptual preparation occurs approximately at 200 ms, followed by word retrieval processes that lead to articulation at 600 to 1200 ms post-picture onset (Indefrey, 2011). However, articulation times are highly variable depending on the nature of the task. Future studies could collect mean reaction times during the overt naming task to pinpoint the onset of covert naming processes.

To conclude, our study highlights the oscillatory patterns associated with the successful integration of newly learned words into existing lexical-semantic networks through learning. We hypothesize that our findings showing a pattern of alpha-beta desynchronization during the covert naming of learned words reflects retrieval after successful consolidation. This evidence aligns with previous research linking alpha and beta activity to lexical-semantic retrieval in different tasks. Our study also paves the way for future investigation into the neural correlates of word retrieval across different phases of language learning. Although we focused on new word retrieval during the first and fifth day of training, future studies could benefit from measuring retrieval over several days and weeks post-training, to better elucidate the time course of lexicalization.

## Acknowledgements

We thank all the participants for their engagement in the present study. We also thank two anonymous reviewers for their constructive comments on a previous version of the manuscript. This work was supported and co-funded by the BrainTrain Research Center of Excellence at the Åbo Akademi University (2015-2018, grant to M.L.) and the Ministerio de Economía, Industria y Competitividad, through the project PSI2015-69132-P to C.F. and the project PGC2018-099859-B-I00 to A.R.F. Moreover, M.L. was supported by funds from the Academy of Finland (Grants No. 260276 and 323251). P.L.C. was funded by a Margarita Salas postdoctoral grant (Ministerio de Universidades, Spain). A.Z. was funded by the European Union’s H2021-MSCA-IF-2021 (project BraSILL No. 101062671). ARF has been suported by the FIAS fellowship Program, co-funded by the European Commission, Marie-Skłodowska-Curie Actions - COFUND Program (Grant #945408). Finally, this work, carried out within the Institute of Convergence ILCB, was supported by grants from France 2030 (ANR-16-CONV-0002) and the Excellence Initiative of Aix-Marseille University (A*MIDEX)

## Authors contribution

NRE, CF, ML, and ARF planned the study. NRE performed the EEG measurements. PLC analyzed the data and performed the statistical analyses. AZ, PLC, CF, ML, and ARF contributed to the interpretation of the data and wrote the manuscript.

## Data and code availability statement

The anonymized data and Matlab scripts necessary to reproduce the analyses presented here are available on the OSF platform (https://osf.io/jzwb2). The materials necessary to attempt to replicate the findings presented here are not publicly accessible but are available from the corresponding author upon request.

