## Supplementary materials for "Alpha and beta desynchronization during consolidation of newly learned words"

**Supplementary analysis: analysis of incorrect trials**

We further investigated the link between alpha-beta power decreases and successful learning and retrieval. With this aim, we conducted an exploratory analysis examining oscillatory changes between Day 1 and Day 5 on incorrect trials corresponding to items not uttered or incorrectly produced in the last overt naming task on Day 5. Note, however, that incorrect trials did not necessarily correspond to “unlearned” items since some were produced in the overt naming task, but not in the correct form (i.e., a phoneme or syllable was wrong). Nonetheless, these trials are still informative as they correspond to items that did not reach the same degree of retrieval as those classified as correct (i.e., correctly produced).

A key limitation in this analysis is the reduced number of incorrect trials available per participant. Indeed, the number of incorrect trials per participant was significantly lower than that of correct items, both on Day 1, *t*(18) = 4.56, *p* < .001 (correct trials, M = 56.7, SD = 13.4; incorrect trials, M = 30.4, SD = 13.9), and on Day 5, *t*(18) = 4.43, *p* < .001 (correct trials, M = 57.3, SD = 14.7; incorrect trials, M = 30.5, SD = 15). This limitation compromises the comparability between correct and incorrect trial results, as null effects observed for incorrect trials may stem from reduced statistical power rather than a true absence of an effect. To control for this confound, we conducted analyses on both trial types (correct and incorrect), ensuring a balanced number of participants and trials across conditions. Furthermore, we controlled for potential stimulus repetition effects by equating trial numbers. Given an equal number of trials across conditions, any effects associated with stimulus repetition should be present in both conditions.

**Participants**. To make the analysis feasible, given the low number of incorrect trials per participant, we softened the exclusion criteria used in the main analysis (see section *2.1 Participants*): we established that participants must have at least 20 available trials (instead of 35 trials) in each condition (correct and incorrect). Please remember that correct trials were excluded if they were already learned on Day 1 (see section *2.1 Participants*). After applying this criterion, a total of 10 participants were included in the analyses.

**EEG analyses**. The same preprocessing pipeline used for correct trials was applied to the incorrect trials (see section 2.5. EEG data acquisition and analyses). After artifact rejection, a total of 10 participants fulfilled the criteria for inclusion in the final analyses.

Next, we balanced the number of correct and incorrect trials. For each participant and session (Day 1 and Day 5), all trials from the condition with fewer available items were retained, and an equal number of trials was randomly selected from the other condition. For instance, if a participant had 45 correct trials and 20 incorrect trials on Day 1, then all incorrect trials (i.e., 20 trials) were included in the analysis, and we randomly drew 20 trials from the pool of 45 correct trials.

As a result, participants had an average of 32 trials in each condition (correct and incorrect) on Day 1 (SD = 7.24) and of 32.9 trials (SD = 7.70) on Day 5. There were no statistically significant differences between the average number of trials on each session (Day 1 and Day 5), t(9) = 0.52, p = 0.62.

**Time-frequency analyses.** All the statistical analyses below were performed using the same parameters in the main analysis (see section *2.6. Time-frequency analyses*). In short, time-resolved power was computed by applying a Hanning taper with a fixed length of 300 ms from 1 to 40 Hz in steps of 20 ms in the -1.1 to 2 s time-window, time-locked to the onset of the picture. For statistical analyses, the grand-averaged time-frequency data were normalized to the 100-ms pre-stimulus period, and a cluster-based permutation analysis was applied on the -0.1 to 1-second interval time-locked to picture onset, including all electrodes and frequencies from 2 to 35 Hz. Specifically, every sample (frequency x time x channel) was compared between the two sessions (Day 1 and Day 5) using a paired-sample-statistic. All samples that exceeded the t-value threshold of ± 2.10 (for an alpha level of 0.05 with 18 degrees of freedom) were selected and clustered based on spectral, temporal, and spatial adjacency. The t-values within every cluster were summed, and the maximum cluster-level sum was used to compute the permutation p-value through the Monte Carlo method involving the maximum number of unique random permutations.

**Time-frequency results**.

***Incorrect trials.*** **Supplementary Figure 1** shows the grand-averaged time-frequency maps of the covert naming task in each session (Day 1 and Day 5) for incorrect trials. For comparability, the electrodes used in the plots are the same as those plotted for the main analysis on the correct trials (see section 3.2. Time Frequency Results). Visual inspection suggests that, on Day 5, picture presentation might have induced a stronger power increase in the upper beta band (25-35 Hz) earlier in the time window and a power decrease within the alpha and beta range (8-25 Hz) in a later interval (**Supplementary Figure 2**), relative to Day 1. However, crucially, there were no statistically significant differences between sessions (Day 1 vs. Day 5) (*p* < .05 on all clusters; the lowest *p-value* for negative clusters was *p* = .45).

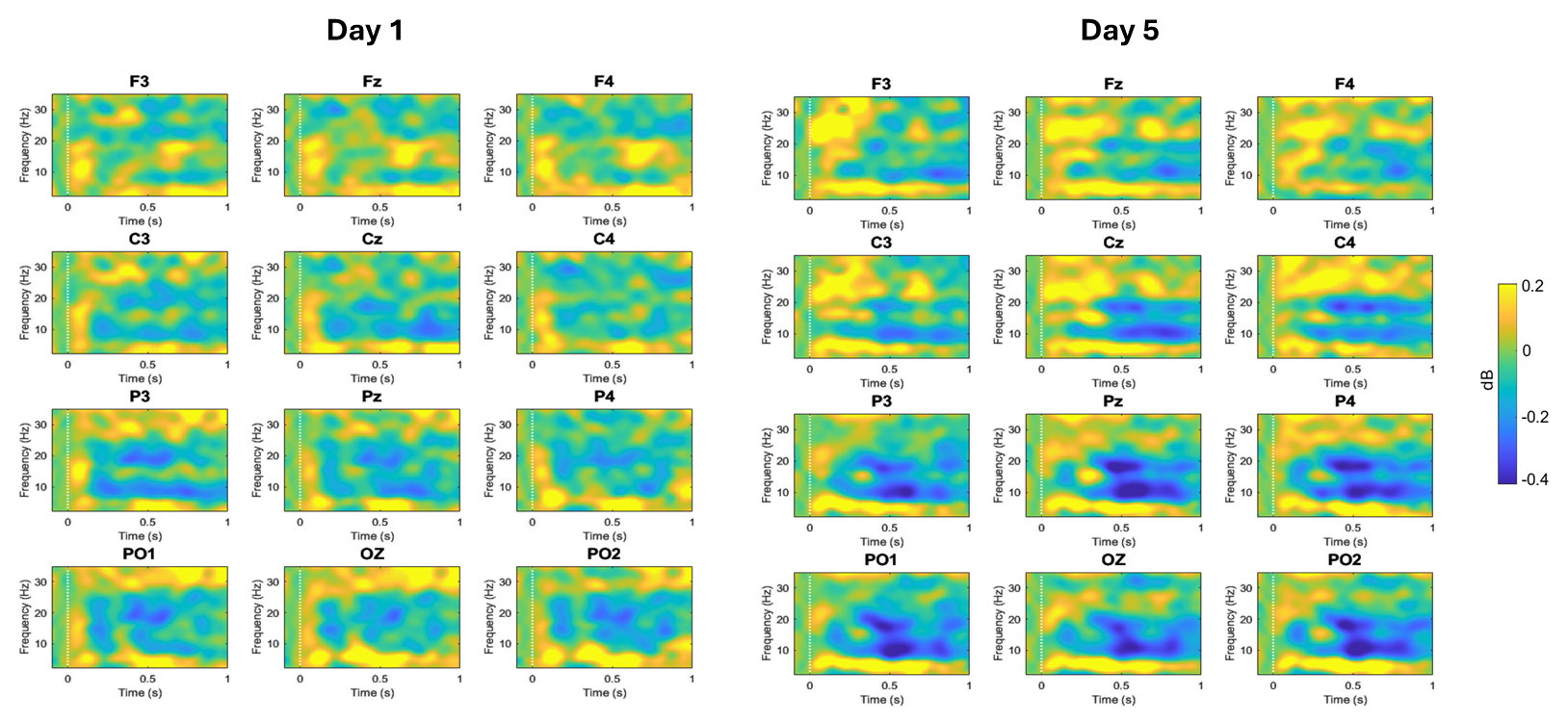

**Supplementary Figure 1**. Time-resolved power for incorrect trials (i.e., not produced or incorrectly produced in the overt naming task) on Day 1 and Day 5 from -0.1 to 1-second post picture-onset and from 2 to 35 Hz. Power ranges from -0.4 to 0.2 dB.

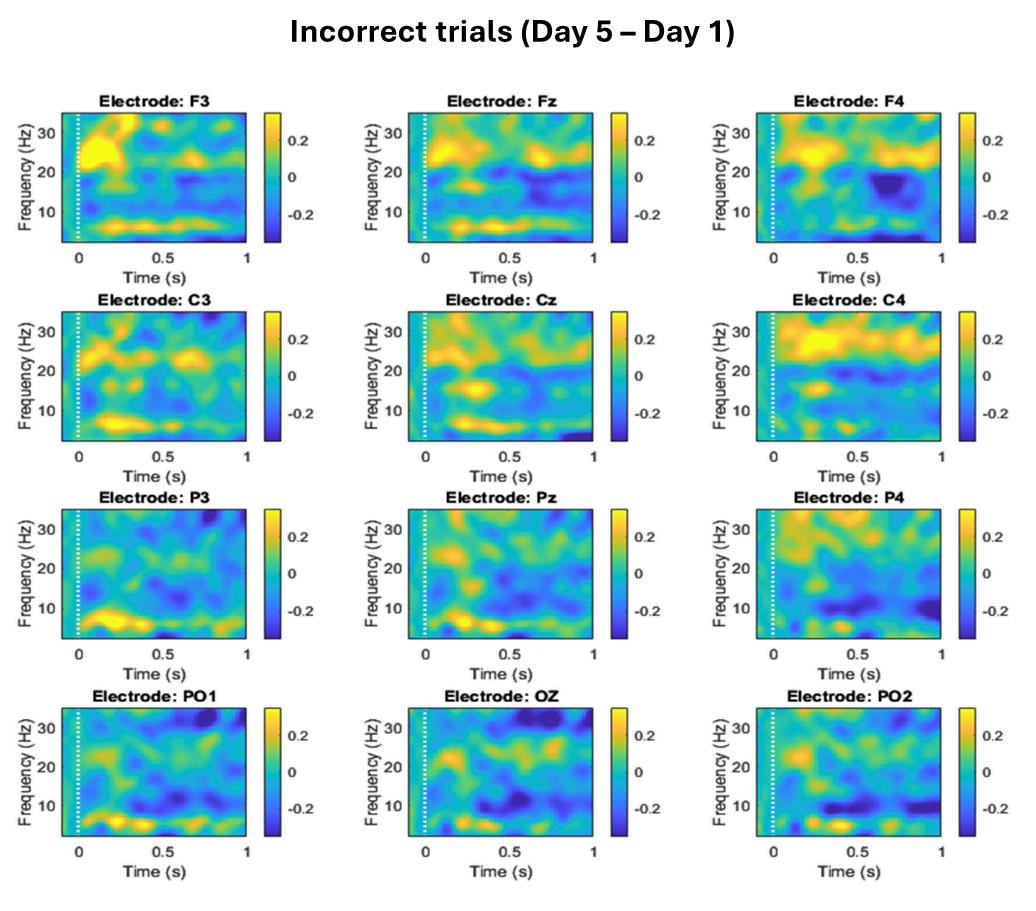

**Supplementary Figure 2.** Time-resolved power changes (Day 5 minus Day 1) for incorrect trials from -0.1 to 1-second post picture-onset and from 2 to 35 Hz. Power ranges from -0.4 to 0.4 dB.

***Correct trials.*** Grand-averaged time-frequency maps of the covert naming task in each session (Day 1 and Day 5) for correct trials are shown in **Supplementary Figure 3**. As observed, picture presentation induced a power decrease within the alpha and beta range (8-25 Hz) on both days, but seemingly more pronounced on Day 5 than on Day 1. Crucially, this was supported by the cluster-based permutation test, which identified a statistically significant negative cluster (*p* < .001), indicating greater power suppression on Day 5 relative to Day 1 (**Supplementary** **Figure 4**). This result aligns with those in the main analysis using a larger sample of participants (N = 19).

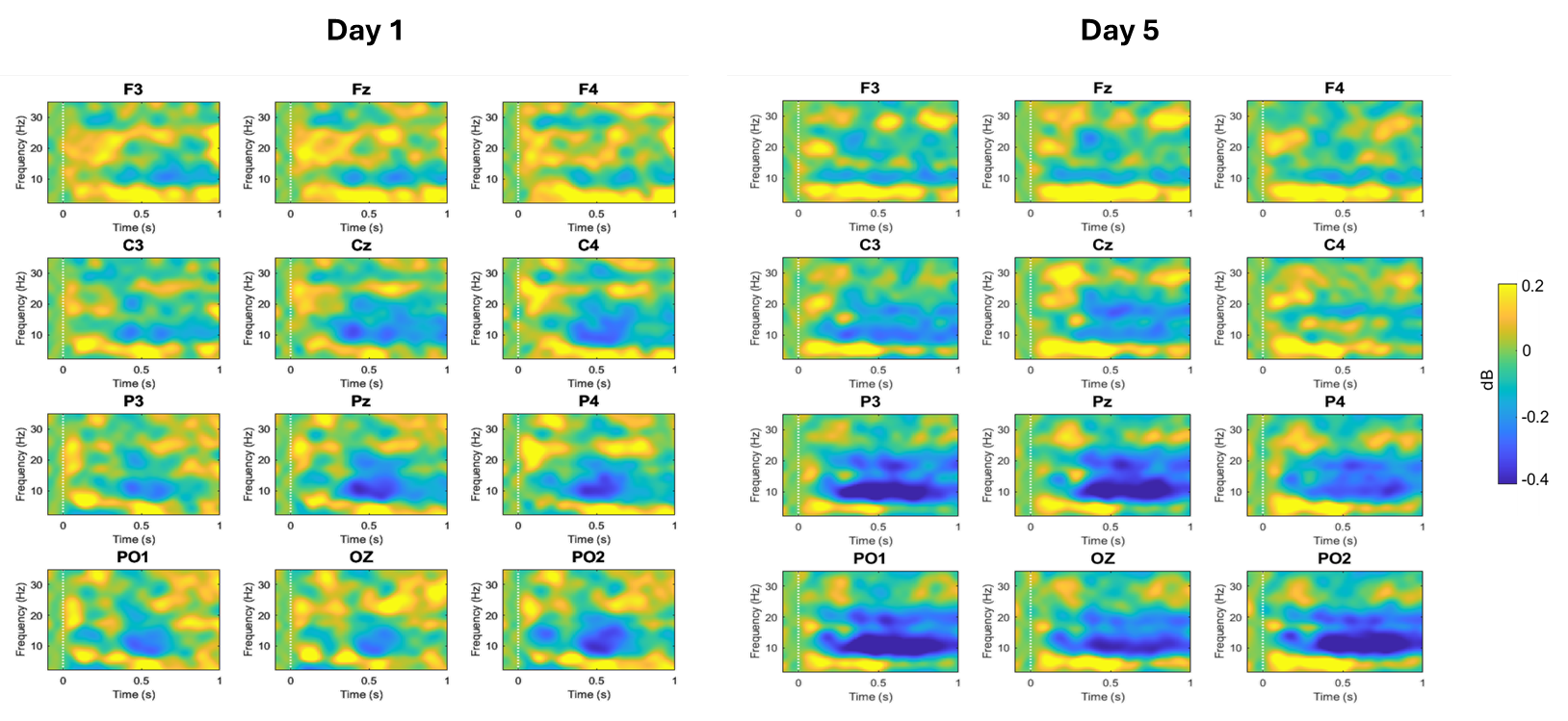

**Supplementary Figure 3**. Time-resolved power for correct trials (i.e., not produced or incorrectly produced in the overt naming task) on Day 1 and Day 5 from -0.1 to 1-second post picture-onset and from 2 to 35 Hz. Power ranges from -0.4 to 0.2 dB.

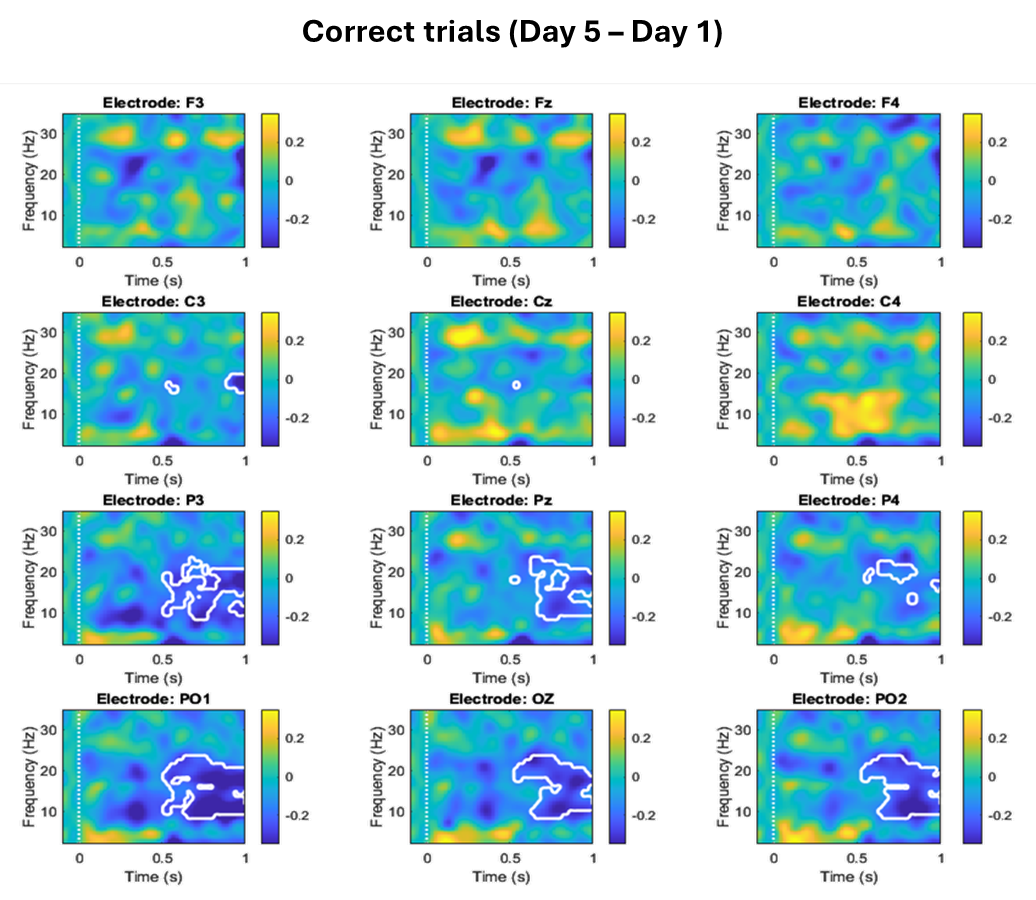

**Supplementary Figure 4.** Time-resolved power changes (Day 5 minus Day 1) for correct trials from -0.1 to 1-second post-picture onset and from 2 to 35 Hz. Power ranges from -0.4 to 0.4 dB.

***Discussion*.** The finding of an effect in correct but not in incorrect trials supports the interpretation that alpha–beta power decrease for correct trials is related to successful consolidation and retrieval, since these trials represent items that were clearly learned and successfully retrieved. Although some desynchronization is also visible for incorrect trials, this may likely reflect partial learning and some degree of retrieval, at least for some items. Thus, by using performance in the overt naming task to distinguish between items that were correctly retrieved (correct) and those that were not, or to a lesser extent, the analysis enables to interpret these neural correlates as reflecting word retrieval after successful consolidation.

Another significant point that emerges from these results is that the alpha–beta power decrease found in correct trials is unlikely attributable to stimulus repetition, given that the number of trials was matched across trial types. If repetition were the main driver, similar effects would be expected in both correct and incorrect trials. Therefore, while repetition may have contributed to some extent, it is not sufficient to account for the observed effect.

Nevertheless, these findings should be interpreted with caution, especially null effects. Considering the small sample size (n = 10) and number of trials per participant, this analysis was most likely statistically underpowered. A more rigorous control for stimulus repetition would require a within-participant design with items that cannot be learned, as done in previous studies (Elmer et al., 2021, 2022).
